# Decoding Remapped Spatial Information in the Peri-Saccadic Period

**DOI:** 10.1101/2023.11.07.565952

**Authors:** Caoimhe Moran, Philippa A. Johnson, Ayelet N. Landau, Hinze Hogendoorn

## Abstract

It has been suggested that, prior to a saccade, visual neurons predictively respond to stimuli that will fall in their receptive fields after completion of the saccade. This saccadic remapping process is thought to compensate for the shift of the visual world across the retina caused by eye movements. To map the timing of this predictive process in the brain, we recorded neural activity using electroencephalography (EEG) during a saccade task. Participants made saccades between two fixation points while covertly attending to oriented gratings briefly presented at various locations on the screen. Data recorded during trials in which participants maintained fixation were used to train classifiers on stimuli in different positions. Subsequently, data collected during saccade trials were used to test for the presence of remapped stimulus information at the post-saccadic retinotopic location in the peri-saccadic period, providing unique insight into *when* remapped information becomes available. We found that the stimulus could be decoded at the remapped location ∼180 ms post-stimulus onset, but only when the stimulus was presented 100-200 ms before saccade onset. Within this range, we found that the timing of remapping was dictated by stimulus onset rather than saccade onset. We conclude that presenting the stimulus immediately before the saccade allows for optimal integration of the corollary discharge signal with the incoming peripheral visual information, resulting in a remapping of activation to the relevant post-saccadic retinotopic neurons.

**Significance Statement:** Each eye movement leads to a shift of the visual world across the retina, such that the visual input before and after the eye movement do not match. Despite this, we perceive the visual world as stable. A predictive mechanism known as saccadic remapping is thought to contribute to this stability. We use a saccade task with time-resolved EEG decoding to obtain a fine-grained analysis of the temporal dynamics of the saccadic remapping process. Probing different stimulus-saccade latencies and an array of stimulus locations, we identify when remapped information becomes available in the visual cortex. We describe a critical window in which feedforward visual information and the preparatory motor signals interact to allow for predictive remapping of a stimulus.

## 1 Introduction

Saccadic eye movements lead to shifts of the visual world across the retina, such that a visual object is represented by different populations of visual neurons before and after the saccade. Despite this, we perceive a stable visual world, undisrupted by intervening saccades. This *spatial constancy* is thought to be achieved through the combination of predictive information about the impending saccade and sensory information from the visual system (Von Helmholtz, 1867; Hallet & Lightstone, 1976; Bays & Husain, 2007; Medendorp, 2011). In this interpretation, just before the saccade, visual neurons rapidly shift or “remap” visual stimuli from their current retinotopic position to the future retinotopic position (i.e. their position on the retina after a saccade). It can be thought of as a predictive ‘glance’ at the post-saccadic retinotopic scene before the eyes leave the current fixation.

The generally accepted explanation of predictive remapping is that, when a saccade is initiated, an internal copy of the motor command, termed the corollary discharge, is sent to sensory systems. This contains information about the direction and amplitude of the upcoming saccade, so that the change in sensory input can be predictively anticipated (Sommer & Wurtz, 2002). Predictive remapping has been demonstrated in a range of experimental paradigms. Early research used single unit recordings in monkeys to shed light on the predictive processes underwriting active vision. Researchers found that during saccade planning, which occurs 200 ms prior to saccade onset (Fischer & Ramsperger, 1984), a single neuron would start to fire in response to a stimulus presented outside of its spatial receptive field (RF) if the upcoming saccade would shift that stimulus into its RF (Duhamel, Colby & Goldberg, 1992). This has been observed in numerous brain areas including visual areas V2, V3 and V3A (Nakamura & Colby, 2002), the lateral intraparietal area (LIP) (Duhamel et al., 1992; Gottlieb, Kusunoki & Goldberg, 1998; Kusunoki, Gottlieb & Goldberg, 2000), frontal eye fields (Bruce & Goldberg, 1985; Thompson, Hanes, Bichot & Schall, 1996; Umeno & Goldberg, 2001) and the superior colliculus (Walker, Fitzgibbon & Goldberg, 1995; Mays & Sparks, 1980). This predictive neuronal firing has been interpreted as a shift of neuronal receptive fields prior to saccade onset, allowing stabilisation of vision despite perturbations caused by the moving eye.

The evidence for remapping in humans has mainly come from psychophysical studies and is often characterised as an updating of attention around the time of a saccade. Cavanagh, Hunt, Afraz and Rolfs, (2010) proposed that the phenomenon of receptive field remapping, as described in animal neurophysiological findings, may be explained by the transfer of attentional states between neurons. The ability of the visual system to remap attention prior to a saccade allows for attentional allocation at the relevant locations, despite disruptive saccades (Marino and Mazer, 2016). Using a double-step saccade task, Rolfs, Jonikaitis, Deubel and Cavanagh (2011) found anticipatory discrimination facilitation at a stimulus’ remapped location prior to a saccade. In other studies, attention has been cued during the pre-saccadic period, showing that attention predictively shifts to the remapped location of the cue in anticipation of the saccade landing (Szinte, Jonikaitis, Rangelov & Deubel, 2018; Jonikaitis, Szinte, Rolfs & Cavanagh, 2013). For example, Szinte et al. (2018) found that participants had increased sensitivity to an orientation change at the remapped location shortly before saccade onset. Similar predictive shifts of attention have been found using a visual masking paradigm, ultimately showing that when a mask is presented at the post-saccadic retinotopic position of a target, discrimination performance is significantly reduced. This is explained by a shift of attention to the mask location leading to interference with perception of the stimulus (Hunt & Cavanagh, 2011). Although these studies have been instrumental in demonstrating attentional updating in the peri-saccadic period, to what extent spatially-specific stimulus information is transferred to the post-saccadic retinotopic location is still up for debate.

In contrast to the direct demonstration of predictive remapping in animal neurophysiological recordings, neurophysiological evidence for predictive remapping in humans is less clear. Predictive remapping involves a precise shift in spatial positions that occurs only briefly, tightly synchronised to saccade execution. Fully characterising this process requires greater concurrent spatial and temporal precision than has been offered by conventional non-invasive human neuroimaging approaches. For example, functional magnetic resonance imaging (fMRI) studies, employing cross-hemispheric paradigms, have been a popular choice in investigating remapping (Medendorp, Goltz & Vilis, 2005; Merriam, Genovese & Colby, 2003, 2007). In such paradigms, with each saccade, the visual stimulus crosses the vertical meridian, which means that the stimulus must be remapped from the current contralateral hemisphere to the future contralateral hemisphere (Parks & Corballis, 2010). Several studies have found that a stimulus flashed in the pre-saccadic period elicits activation in the ipsilateral hemisphere which was never directly stimulated (Merriam et al., 2003, 2007; Parks & Corballis, 2010). However, the temporal resolution of fMRI is limited, allowing only a very coarse insight into the time course of neural computations. Understanding the temporal dynamics of the saccadic remapping process is essential to understanding the underlying mechanisms; establishing when the brain has access to relevant visual information, whether it be before (pre-saccadic), during (trans-saccadic) or only after (post-saccadic) the saccade, is necessary to understand how the visual system maintains visual stability.

In order to provide this necessary temporal precision, other studies have attempted to investigate saccadic remapping using electroencephalography (EEG). However, the limited spatial precision of EEG has constrained conclusions about spatial remapping to a very coarse scale. For example, Parks & Corballis (2010) examined event-related potentials (ERPs) and found that if a saccade shifted a stimulus from one hemifield to the other, prior to that saccade, there was an increase in positivity in the hemisphere ipsilateral to the visual stimulus. This shows, at a coarse level, remapping of visual space in anticipation of a saccade. Another study employed a multivariate pattern analysis (MVPA) approach in which participants made a saccade to either a house or a face located in the periphery (Edwards et al., 2018). They reported that when the stimulus remained the same across the saccadic period, they could decode it 123 ms after saccade offset, whereas when the stimulus was changed in the trans-saccadic period, stimulus decoding only became possible later, at 151 ms post-saccade offset. This decoding advantage suggests anticipation of the post-saccadic input before the eyes have moved (Edwards et al., 2018). Importantly, neither of these studies sampled multiple stimulus locations, limiting the conclusions that can be drawn about the spatial specificity of the remapped response. Moreover, both studies involved stimuli that were visible before, during, and after the saccade^1^. Although the interaction between pre-saccadic and post-saccadic input is suggestive of remapping, a direct demonstration of predictive remapping would require evidence that a *pre-saccadic* visual stimulus is represented neurally at its *post-saccadic* retinotopic location, uncorrupted by any input once the eyes start moving. This would ensure that any sensory information about the stimulus was acquired before saccade onset, and therefore that any neural representation of the stimulus in the remapped location is attributable to predictive remapping.

In this study, we applied an EEG decoding approach to investigate the neural representation of stimulus position in the presence of a saccade. Stimuli were briefly presented before saccade onset and removed before saccade landing. This ensured that there was no direct visual stimulation at the remapped location. This novel analysis approach, in combination with a careful experimental design, allowed us to characterise the fine-grained time-course of neural position information for stimuli presented in multiple different positions and at different stimulus-saccade latencies. Using this approach, we show that visual stimuli presented during a narrow temporal window 100-200 ms before a saccade, are subsequently represented in the visual cortex in their future post-saccadic retinotopic positions, providing strong neural evidence of predictive remapping in humans.

## 2 Materials and Methods

### 2.1 Participants

Participants were required to complete six testing sessions across different days, including one screening session. The screening session provided a sanity check to ensure above-chance decoding of stimulus position on fixation trials. This was used as an inclusion criterion. In Protocol 1, fifteen observers took part in the initial screening. Six participants were excluded after analysis of their screening session data, due to poor EEG classification performance (less than 52% average decoding accuracy when classifying stimulus presentation location on fixation trials), and an additional participant was excluded after completing all six sessions due to data collection errors. After exclusion, eight observers (four male; mean age = 26.1 years, sd = 5.5 years) with normal or corrected-to-normal vision remained. In Protocol 2, thirteen observers took part in the initial screening session, of whom three were excluded due to poor EEG classification performance (less than 52% average decoding accuracy when classifying stimulus presentation location on fixation trials). After exclusion, ten observers (three male; mean age = 25.7 years, sd = 2.0 years) with normal or corrected-to-normal vision remained. Participant data was combined across the protocols giving a total of 18 participants (seven male; mean age = 26.4 years, sd = 4.1 years). The experimental protocols were approved by the human research ethics committee of The University of Melbourne, Australia (Ethics ID: 2021-12985-16726-4) and conducted in accordance with the Declaration of Helsinki. All observers provided informed consent before beginning the experiment and were reimbursed AU$15 per hour for their time, plus an additional AU$20 after completing all sessions.

### 2.2 Stimuli and procedure

Stimuli were programmed in MATLAB Version R2020a, using the Psychophysics Toolbox extension (Brainard, 1997; Pelli, 1997; Kleiner, Brainard & Pelli, 2007). They were presented on an ASUS ROG PG258 monitor (ASUS, Tapei, Taiwan) with a resolution of 1,920 × 1,080 running at a refresh rate of 120 Hz. Participants were seated in a quiet, dark room with their head supported by a chin rest, positioned 80 cm from the screen.

Stimuli differed slightly across the two protocols of the experiment. In both protocols, stimuli consisted of sinusoidal gratings presented within a circular Gaussian window (i.e. Gabors; outer diameter: 7.9 degrees of visual angle (dva); 100% contrast) presented on a grey background for 100 ms. In Protocol 1, the stimulus was always presented at a spatial frequency of .16 c/dva (except on catch trials), while in Protocol 2 the stimulus could appear at a spatial frequency of .33 c/dva or 1.13 c/dva. In Protocol 1, grating orientations were evenly spaced between 0° and 150° in 30° steps, meaning six possible orientations could be presented (stimuli were collapsed across orientations for analyses). In Protocol 2, orientation of the grating was fixed at 0° (except on catch trials). For catch trials, participants were instructed to report an oddball grating of either a higher spatial frequency; .51 c/dva, (Protocol 1) or of a different orientation; 90° (Protocol 2) which could appear every 11-20 trials. These trials were included in order to ensure attentive processing of the grating stimuli, and were excluded from the main analysis. All other details were the same across the two experimental protocols.

Two fixation points (.41 dva), one black and one white, were horizontally aligned and subtended 10.04 dva to the left and right of the screen centre (20.08 dva apart). They appeared at the beginning of the experiment and remained visible throughout, excluding breaks. Participants were instructed to fixate on the black fixation point at all times. Periodically during the experiment, a saccade cue appeared: the colour of the fixation points gradually changed, such that over a period of 1.2s the black fixation point became white and vice versa. Participants were instructed to monitor the colour of the fixated fixation point, and plan and execute a saccade from one fixation point to the other as soon as they detected the colour change, such that they were always fixating the black fixation point.

Different conditions are determined by the conjunction of fixation position and the location of the grating. In fixation trials, the fixation is stable (no colour change occurs), and stimuli appear in one of four positions around the current black fixation point (location 1-4; Fig. 1a). In control trials the fixation is stable, and stimuli appear on the other side of the screen, in one of four locations around the white fixation point (location 5-8; Fig.1a). In saccade trials, there is a colour change and stimuli appear during the planning of a saccade. In this case, stimuli can appear around current fixation (not analysed in the present manuscript) or the saccade target (saccade trials) (location 5-8; Fig.1a) (see Fig. 1 b and c for task design).

**Figure 1:**
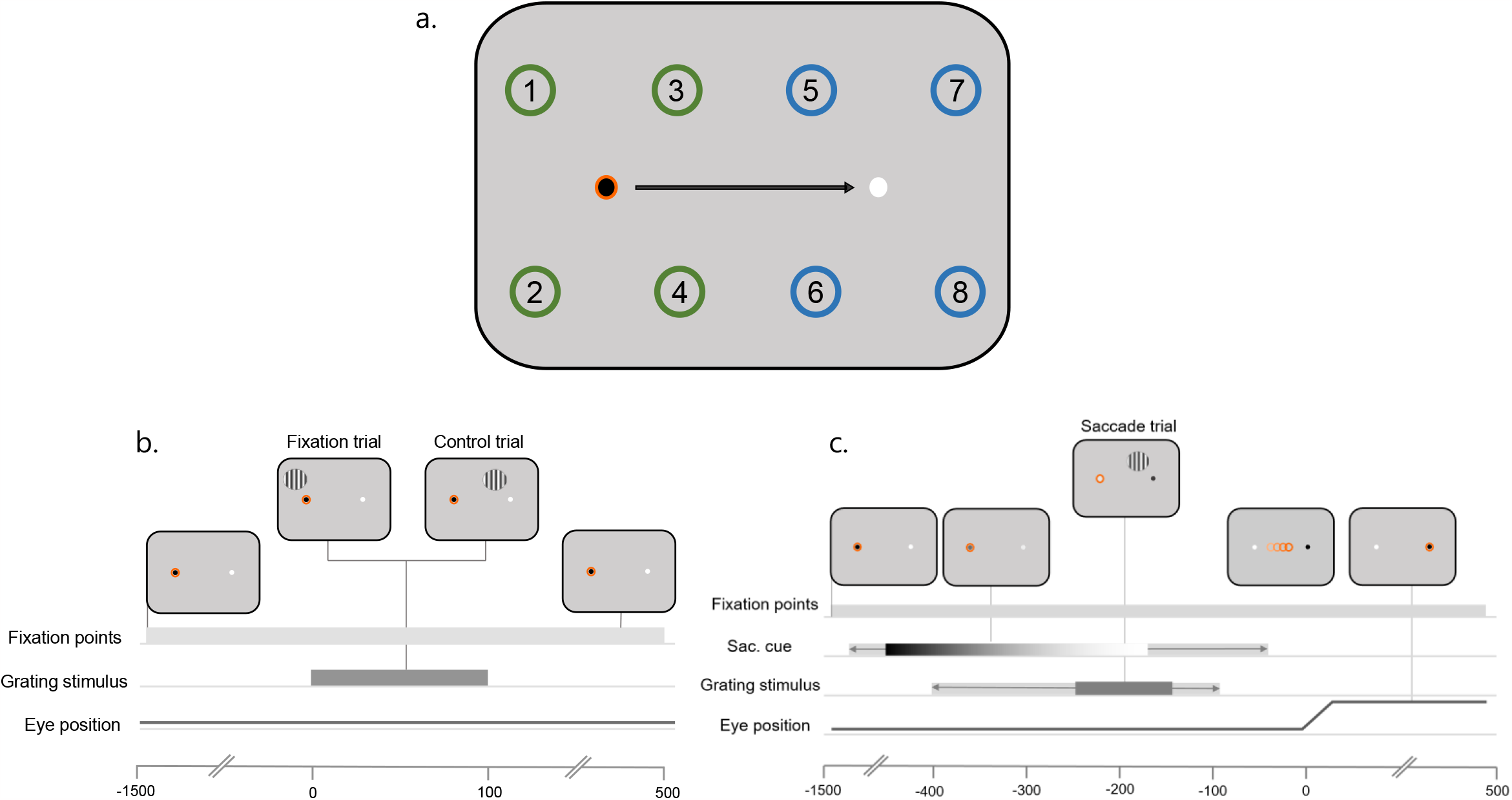
**a. Stimulus configuration.** Participants always fixated on the black fixation point and made a saccade when cued by a gradual colour change to white. Stimuli were presented around fixation on fixation trials (location 1-4) and around the white fixation point on control trials or the saccade target on saccade trials (location 5-8). The orange circle around the black fixation point indicates eye position. The numbers on the screen indicate all possible stimulus presentation locations. The green and blue circles indicate where stimuli could be presented on fixation trials and control or saccade trials, respectively. The black arrow shows saccade direction. All are for illustration purposes only and were not presented to participants. **b**. In **fixation trials**, participants maintained fixation on the black fixation point throughout the trial, never fixating the white. The orange ring indicates the position of the eyes. Approximately 400-800 ms after trial onset, the grating stimulus was presented at one of four locations around fixation. The stimulus was displayed for 100 ms. In **control trials**, participants fixated on the black fixation point. A grating was presented at one of four locations around the alternate fixation point (white circle). **c**. In **saccade trials**, participants fixated on the black fixation point. After 200-600 ms the black fixation point changed to white and the white fixation point changed to black, which served as the saccade cue. The colours gradually changed over a period of 1.2 s, indicated here by the colour gradient. The light grey bar behind the gradient refers to possible start/end times of the colour change. In a period ranging from 400 ms to 100 ms before saccade onset, a grating was presented at one of four locations around the saccade target. The stimulus was presented for 100 ms. The light grey bar behind dark grey bar refers to possible start/end times of the stimulus.

Stimulus locations were separated by 5.06 dva and were also 5.06 dva away from the nearest fixation point (Fig. 1a). The grating was presented for 100 ms at a variable delay after the saccade cue. This delay was adjusted for each participant in order to sample trials with approximately 200 ms between stimulus onset and saccade onset. To do so, the latency between grating onset and saccade onset was recorded on every trial and averaged over the previous 100 trials (see Fig. 4 for the distribution of stimulus-saccade onset latencies).

Each experimental session contained a total of 2,400 trials: 1,120 fixation trials, 1,120 saccade trials and 160 control trials, randomly interleaved across 5 blocks. There were a total of 480 trials per block, with a mini-break every 100 trials. Trials were split evenly between fixation and saccade trials (224 trials each) with the remaining allocated to control trials (32 trials).

### 2.3 EEG and eye-tracking pre-processing

EEG and EOG data were recorded at 2048 Hz using a BioSemi system, with 64 active electrodes and 6 ocular electrodes. The continuous EEG and eye-tracking data was pre-processed off-line using MATLAB Version R2020a and EEGLAB toolbox (v2021.0) (Delorme & Makeig, 2004). The data was first down-sampled to 256 Hz and then re-referenced to the mastoids. Eyetracking data were recorded using an Eyelink 1000 eyetracker (SR Research) at 1000 Hz. The eyetracker was calibrated at the start of the experiment and at the beginning of every block. In addition, drift correction was applied at each mini break within a block. The eye-tracking data was synchronised with the EEG using the EYE-EEG toolbox version 0.81 (Dimigen, Sommer, Hohlfield, Jacobs & Kliegl, 2011b). The EEG data was then notch filtered at ∼50Hz to remove electrical artifacts and bandpass filtered between 0.1 and 80 Hz. Automatic data rejection was employed to remove any major artifacts using the Artifact Subspace Reconstruction (ASR) method in EEGLAB. The ASR Rejection threshold Parameter *k* was set to 15. Bad channels, noted during data collection and confirmed later offline, were spherically interpolated.

To correct for eye movement artifacts in the EEG, we applied independent component analysis (ICA; (Makeig, Jung, Ghahremani & Sejnowski, 1996)). To identify eye movement related components, the variance ratio of the component activation during periods of eye movements (blinks and saccades) was compared to that during fixation periods (Plöchl, Ossandón, & König, 2012). ICA was performed in a separate pre-processing pipeline containing an additional high-pass filter (Hamming windowed sinc FIR, edge of the passband: 2Hz). The ICA weights were then appended to the corresponding datasets. Components were rejected if the mean variance of activity selected near a saccade was 10% greater than the mean variance during fixation periods (Dimigen, 2020; Plöchl et al., 2012). When gaze deviated more than 2.16 dva from the fixation point or landed more than 2.16 dva away from the saccade target, the trial was discarded. On average, 5% of saccade trials were removed per subject.

EEG data were epoched for fixation, control and saccade trials separately. Across all trial types, epochs were time-locked to the presentation of the grating. Epochs were extracted from 200 ms before stimulus onset to 500 ms after and were baseline corrected to the mean of the 100 ms period before stimulus onset. Fixation trials were subsequently included in the training set for the main analysis.

### 2.4 Multivariate Pattern Analysis

Epochs from the training set of fixation trials were used to train time-resolved pairwise linear discriminant analysis (LDA) classifiers (Grootswagers et al., 2017) to dissociate the neural activation patterns associated with the presentation of the stimulus at two different positions, using the amplitude from the 64 electrode channels as features. This set of LDA classifiers was tested on each pairwise combination of the four possible stimulus presentation locations on independent fixation trials and subsequently averaged across all pairs. This was done separately for left and right fixation trials, across all timepoints in the epoch and for each individual participant. An above-chance classification performance indicates that the EEG signal contained information that allowed the classifier to distinguish a stimulus presented at one location versus another. The same trained classifiers were used to decode stimulus location on control trials and saccade trials.

To examine the time course of remapped stimulus location information, the classifiers that were trained to dissociate stimulus locations on fixation trials were tested using data from the peri-saccadic period of saccade trials. time-locked to stimulus onset. As an initial step, trial-by-trial classification time courses were split into three bins depending on when the saccade occurred relative to stimulus onset (i.e. stimulus-saccade onset asynchrony; SSOA), for the purposes of visualisation and statistical inference. We sorted saccades into a Short Bin (SSOA = 100-200 ms), a Medium Bin (200-300 ms) and a Long Bin (300-400 ms). ‘Short’, ‘medium’ and ‘long’ refer to the time between stimulus onset and saccade onset. In order to probe the precise timing of remapping we then used a more fine-grained analysis in which a sliding window, with a window length of 100 ms and a step size of 10 ms, was moved along the saccade trial time course.

### 2.5 Statistical inference

We used Bayes factors (BFs) to determine above- and below-chance decoding and at chance decoding (i.e., null hypothesis) at every timepoint within each of the eighteen participants using the Bayes Factor R package (Morey & Rouder, 2018) implemented in Python (Teichmann, Moerel, Baker & Grootswager, 2021). We set the prior for the null hypothesis at 0.5 (chance decoding) for assessing the decoding results and at 0 for assessing differences across decoding results. A half-Cauchy prior was used for the alternative hypothesis with a medium width of r = √22 = 0.707. Based on Teichmann et al., (2021), we set the standardized effect sizes expected to occur under the alternative hypothesis in a range between -∞ and ∞ to capture above and below chance decoding with a medium effect size (Morey & Rouder, 2018).

BFs larger than 1 indicate that there is more evidence for the alternative hypothesis than the null hypothesis (Dienes, 2011). We used a BF cut-off of 10 which is considered substantial evidence for the alternative hypothesis (Wetzels et al., 2011).

### 2.6 Task performance

All participants performed well at the oddball task with a mean accuracy of 98% (SE = 2%) in Protocol 1 and 84% (SE = 7%) in Protocol 2. Reaction times (RTs) were calculated from the time that the oddball stimulus appeared on the screen until the time that the space bar was pressed. The mean RT was 670 ms (SE = 128 ms) in protocol one and 623 ms (SE = 150 ms) in protocol 2.

## 3 Results

### 3.1 Decoding the position of central stimuli

In order to assess whether we could extract location information from the EEG signal, we trained classifiers on a subset of EEG epochs in which stimuli were presented in the four locations around current fixation and tested classifier accuracy using left out fixation trials (Fig. 2*a*). Using a 5-fold cross-validation procedure and pairwise comparisons between stimulus locations, we found that classifiers effectively labelled the test set trials in their correct location across all pairs (Fig. 2*c*). The pairs consisted of diagonal comparisons (i.e. 1 vs 4 and 2 vs 3 in Fig. 1*a*), vertical comparisons (i.e. 1 vs 2 and 3 vs 4; Fig. 1*a*) and horizontal comparisons (i.e. 1 vs 3 and 2 vs 4; Fig. 1a). Overall, performance was very similar across the different comparisons, with vertical, diagonal, and horizontal comparisons peaking at 72%, 67%, and 69% accuracy, at latencies of 164 ms, 160 ms, and 164 ms, respectively (Fig. 2*c*). Together, these results demonstrate that multivariate classifiers were able to extract position information from the EEG signal.

**Figure 2.**
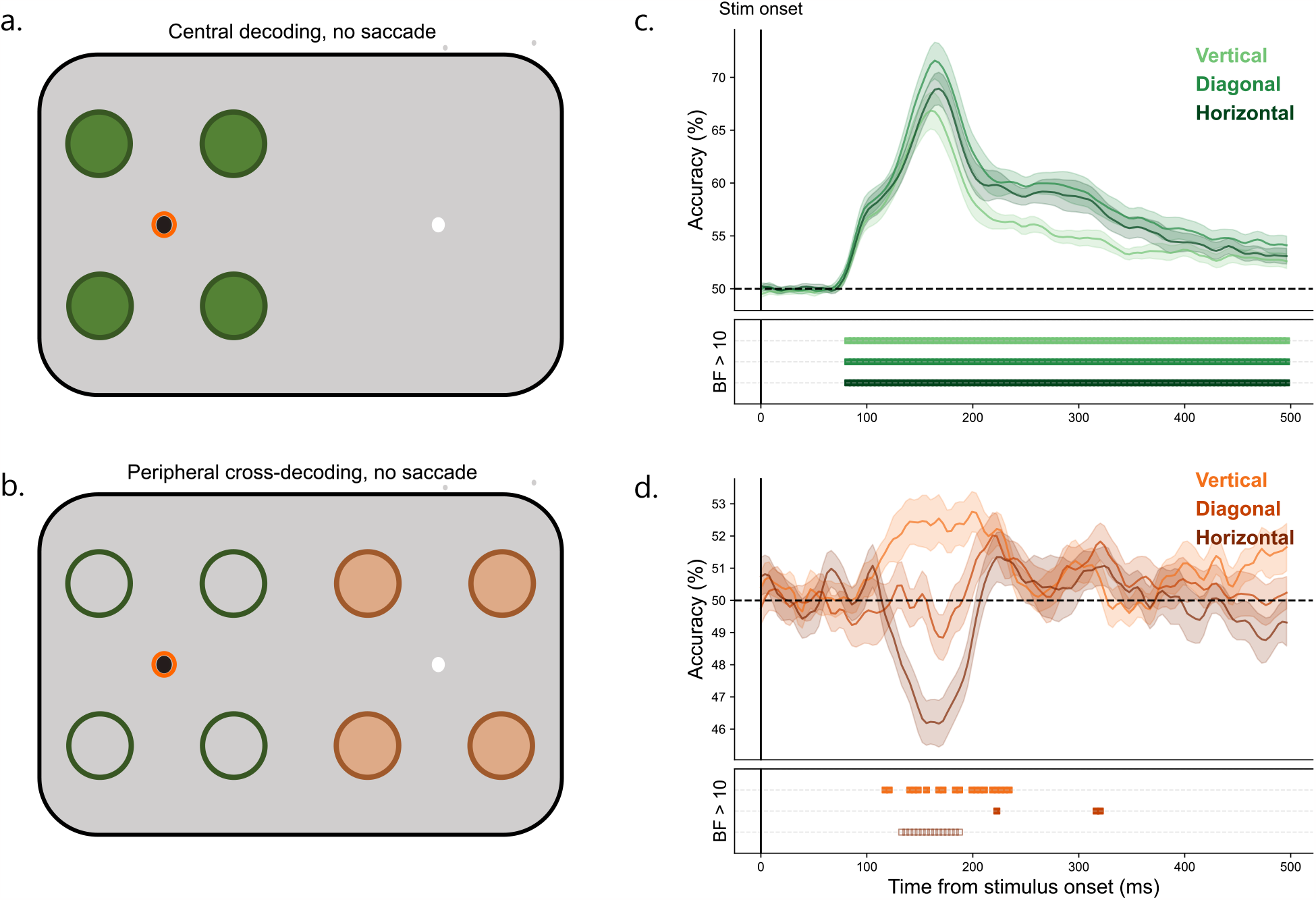
Decoding results across fixation and control trials. This figure illustrates how the classifiers were trained and tested to distinguish neural patterns evoked by the presentation of a stimulus at different locations in **a**. fixation trials and **b**. control trials. In a and b, filled circles represent possible stimulus presentation locations and thus, what the classifiers were tested on. In a, these also correspond with the training locations. The unfilled circles in b indicate the locations that the classifier was trained on (using fixation data). The bright orange ring indicates current fixation. **c**. The mean classification over time for fixation trials. The plotted results reflect the average performance at corresponding training and testing timepoints. The timepoint of peak performance for each subject was subsequently used as training data to test control and saccade trials. **d**. The mean classification over time for control trials. The plotted results reflect the average performance of classifiers trained on the peak time point of fixation trials and tested on the entire control trial time course. Shaded areas depict standard error of the mean across subjects. All decoding results are averaged over the relevant pairwise comparisons, across left and right fixation, and subsequently, across subjects. The Bayes factors (BF) below the plots indicate the timepoints at which there was substantial evidence in favour of the alternative hypothesis i.e. decoding above- (filled squares) or below- (open squares) chance level.

### 3.2 Generalising position decoding to peripheral stimuli

After establishing that classifiers were able to decode the position of a stimulus on fixation trials, we investigated whether position information could be decoded from peripheral stimuli in the absence of a saccade (Fig. 2*b*). To do so, we tested these same classifiers on control trials, in which the stimulus was presented in the periphery but no saccade was made.

Accordingly, the same classifiers that were trained to discriminate pairs of stimulus locations on fixation trials were tested using EEG data collected during control trials, with locations counted as “correct” if the retinotopic training position matched the retinotopic position of the control stimulus, relative to the other fixation point (i.e. 1-5, 2-6, 3-7, 4-8 in Fig. 1a). The timepoint of peak decoding from the trained classifiers (fixation trial data) was tested on every timepoint of control trials (0-500 ms after stimulus onset). This was done on an individual subject level, meaning the timepoint used for training varied across subjects. The mean peak training timepoint was 167 ms (SE = 24 ms) after stimulus onset.

We observed above-chance cross-decoding for both vertical and diagonal pairwise comparisons and below-chance cross-decoding for horizontal pairwise comparisons (Fig. 2*d*). Above-chance decoding in the vertical comparisons indicates that even though classifiers were trained on stimuli near fixation and tested on stimuli in the periphery, central and peripheral stimuli shared EEG features. This is understandable since ERPs evoked by stimuli in the upper and lower visual fields are highly dissociable (Srebro, 1985; Slotnick 2018).

Similarly, diagonal comparisons were above chance for control trials (albeit briefly), due to the upper/lower visual field dissociation. Conversely, we observed below-chance decoding performance for horizontal comparisons. This is most likely attributable to the relative spatial proximity of the mismatched spatial positions. For example, a classifier trained to discriminate between positions 1 and 3 (Fig. 1*a*), and tested on control position 5 is more likely to assign that trial to position 3 than to position 1 due to its relative proximity. This assignment is opposite to the retinotopic locations of the stimuli relative to the two fixation points (i.e. 3 is upper-right in the training set, relative to current fixation, while 5 is upper-left in the test set, relative to the alternate fixation point).

Importantly, because we observed above-chance decoding for control stimuli separated by the horizontal midline (i.e. vertical and diagonal comparisons), only horizontal comparisons were used in subsequent steps to investigate saccadic remapping. This is appropriate because it ensures the most conservative possible approach to demonstrating remapping: without a saccade, decoding performance is significantly *below* chance at the putative remapped location.

### 3.3 Generalising position decoding to peripheral stimuli with a saccade

To investigate whether spatial information is remapped in the peri-saccadic period, we next tested the same fixation classifiers on peripheral stimuli presented during saccade trials (Fig. 3*a*). To gain a better understanding of the unfolding of events from stimulus presentation until after the saccade, we split the saccade trials into three different bins based on stimulus-saccade onset asynchrony (SSOA). The mean saccade latency for the Short Bin (100-200 ms) was 157 ms (sd = 29 ms), for the Middle Bin (200-300 ms) was 254 ms (sd = 28 ms) and for the Long Bin (300-400 ms) was 344 ms (sd = 29 ms) (Fig. 3*b*). The same trained classifier (horizontal comparisons in fixation trial data) were tested on the entire saccade trial (0-500 ms after stimulus onset) across bins. As in the control analysis, classifiers were trained on the timepoint of peak decoding in fixation trials for each participant.

**Figure 3.**
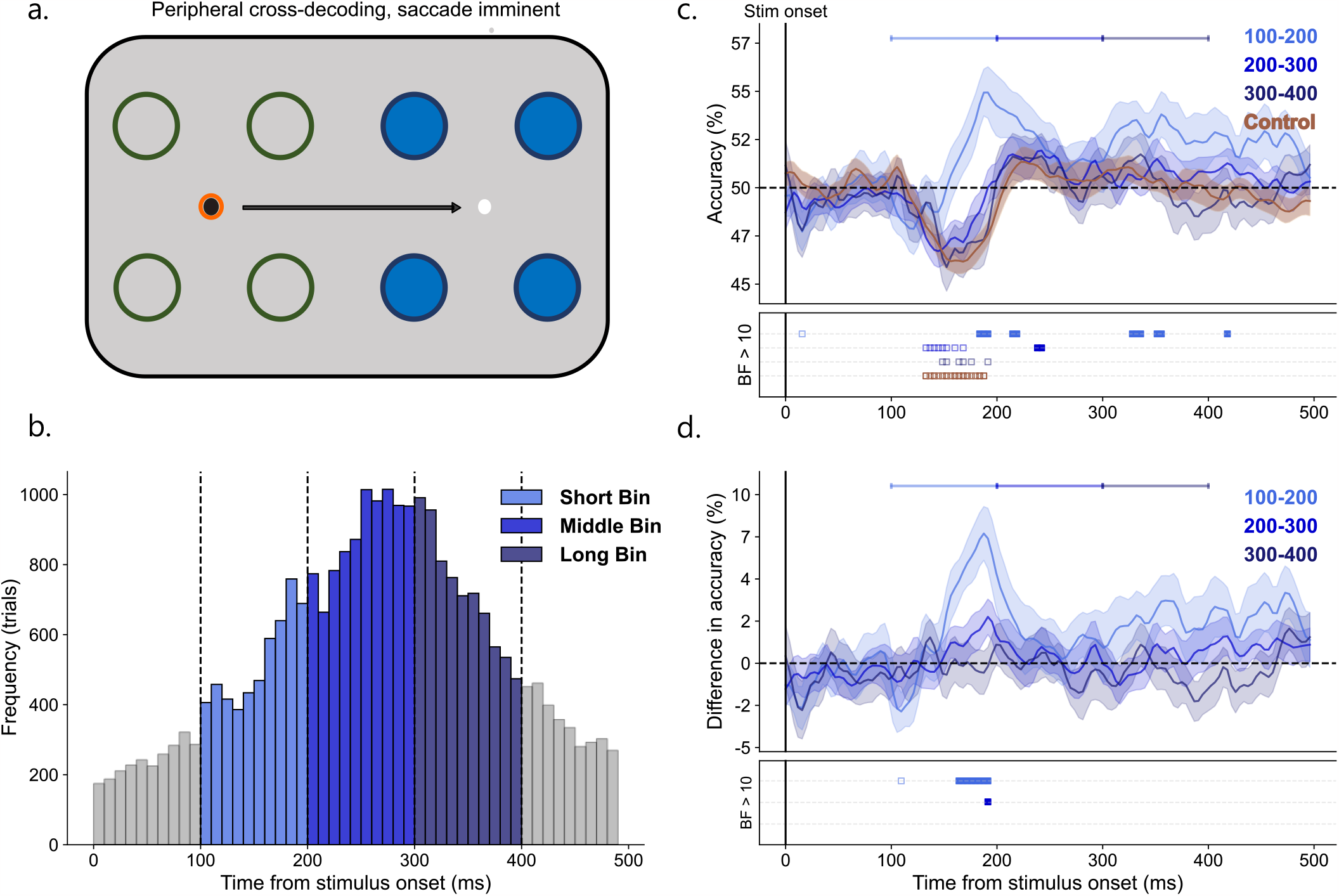
Saccade trial analyses. **a.** Classifier training and testing for saccade trials. The black arrow indicates the planned saccade. Filled circles represent possible stimulus presentation locations and thus, what the classifiers were tested on. The unfilled circles indicate the locations that the classifiers were trained on (using fixation data). The bright orange ring indicates current fixation. **b**. Distribution of saccade latencies relative to stimulus onset are shown, split into three 100 ms bins (Short Bin: 100-200 ms, Middle Bin: 200-300 and Long Bin: 300-400). The grey bins were not considered in this analysis. **c**. Average performance of classifiers trained on the peak time point of fixation trials and tested on control and saccade trials. In this case, classification assignment is considered correct if the stimulus is assigned to the corresponding remapped position around the black fixation point. Different shades of blue represent different saccade latency bins, and brown indicates control trials. The Bayes factors (BF) below the plots indicate the timepoints at which there was substantial evidence in favour of the alternative hypothesis i.e. decoding above- (filled squares) or below- (open squares) chance level. **d)** The difference in classification accuracy between the control condition and all saccade latency bins. Only in the Short Bin were there consistently any timepoints in which there was sufficient evidence of a difference between the saccade condition and the control condition. Dots below plot indicate thresholded Bayes factors (BF) for which there is substantial evidence of a difference in classification accuracy between saccade and control trials. In c. and d., shaded areas depict standard error of the mean across subjects. All decoding results are averaged over the relevant pairwise comparisons, across left and right fixation, and, subsequently, across subjects.

**Figure 4.**
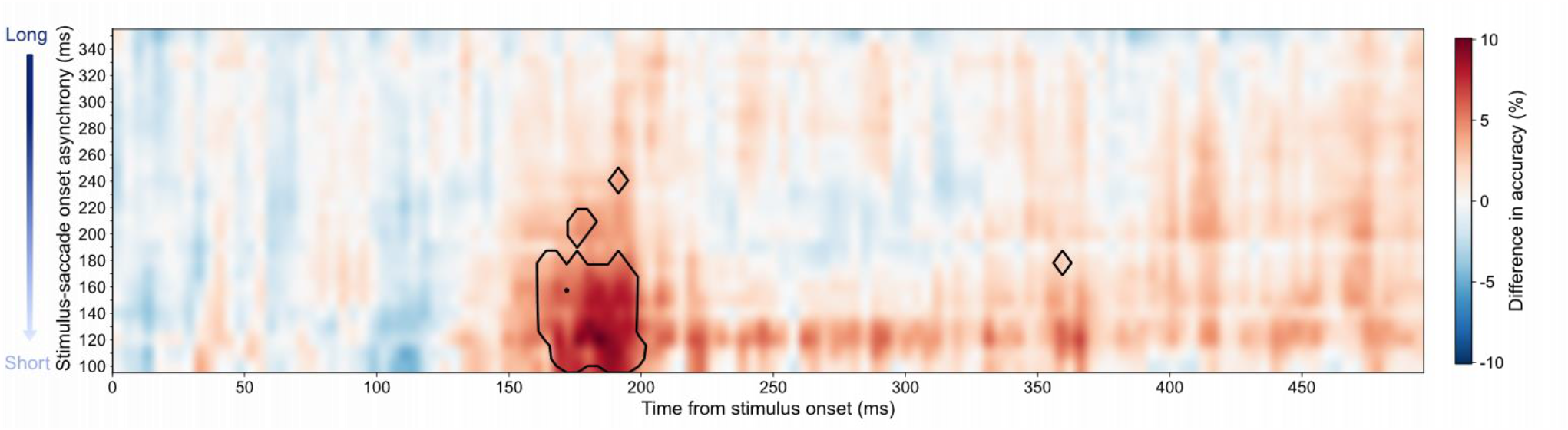
Difference in classification accuracy between control trials and saccade trials. The classification results of control trials were subtracted from each saccade bin to provide accuracy differences. The y-axis shows saccade bin centres ranging from long to short SSOAs i.e. the time between stimulus onset and saccade onset. The classification results for each of the 26 bins occupies a single row in the matrix. The areas outlined in black indicate a BF greater than 10.

Medium and Long latency bins followed a similar pattern of decoding to that seen in control trials, with a below-chance classification peak at 145 ms (47%) and 152 ms (45%), respectively. In contrast, the Short Bin showed the opposite effect, first rising above chance level at 184 ms with peak decoding (55%) at 188 ms post stimulus onset (Fig. 4*c*). We then looked at the difference between decoding accuracy for each of the three saccade bins relative to control trials (Fig. 4*d*). Only the shortest latency bin was consistently and significantly different from control trials, with a peak difference of 9% at 188 ms post-stimulus onset (Fig. 4*d*).

### Fine-grained temporal analysis

Analyses so far have shown that spatial information is remapped for trials in which a stimulus is presented roughly 100-200 ms before a saccade. To further investigate precisely when a saccade must be made for remapping to be evident, and when the remapped information becomes available in the EEG signal, we carried out a further analysis. We again trained classifiers on the peak timepoint of fixation trials (by subject). While saccade trials were previously split into 3 non-overlapping bins, this time we split the trials into overlapping bins, defined using a sliding window that moved from 400 ms to 50 ms before saccade onset. Thus, the bins evolve from long SSOAs to short SSOAs. Each bin was 100 ms long and the window was shifted by 10 ms each time, giving a total of 26 bins (Fig. 4). Sufficient evidence in favour of remapped spatial information (BF > 10) emerged in the early cluster of bins ranging from a bin centre of 100 ms to 180 ms. The largest difference in decoding accuracy (10%) between control and saccade trials occurred in the bin ranging from 70 – 170 ms (bin centre = 120 ms) at 184 ms post-stimulus onset. This indicates that spatial information was remapped only when a saccade occurred approximately 100 - 180 ms after stimulus onset, with strongest evidence for remapping when a saccade occurred ∼120 ms after stimulus onset.

## 4 Discussion

Despite the constant movement of our eyes, we are able to keep track of relevant objects in the world across saccades. This spatial constancy has been attributed to a process known as predictive remapping, in which immediately prior to a saccade, visual neurons respond to stimuli falling in their future, rather than current retinotopic receptive field. Although this mechanism has been extensively demonstrated using invasive neurophysiological recordings in visual and oculomotor areas of primates (Duhamel et al., 1992; Nakamura & Colby, 2002), how predictive remapping manifests in the human brain is less clear. In this study, we used multivariate pattern analysis (MVPA) of EEG data to provide spatiotemporally resolved, neural evidence of saccadic remapping in humans. By examining decoding results under different train-test combinations and a range of stimulus-saccade latencies, we demonstrated that the neural representation of a visual stimulus briefly presented before an impending saccade indeed encodes the predicted future retinotopic position of that stimulus. This remapping was tightly locked to stimulus onset and was most present when the stimulus was shown in a critical period centred on 120 ms before saccade onset. This shows that immediately before a saccade, the visual system predictively represents visual objects in their anticipated future retinotopic positions, which likely plays a key role in maintaining visual stability across eye movements.

The importance of stimulus timing relative to saccade onset suggests that corollary discharge (CD) is the driving force behind this predictive process. We found above-chance decoding of the remapped location only when the stimulus was presented in a narrow window centred on 120 ms before saccade onset. This is when saccade preparation occurs and, by consequence, when the CD signal is available (Wurtz, 2008; Crapse & Sommer, 2008; Cavanaugh, Berman, Joiner & Wurtz, 2016; Sommer & Wurtz, 2008). Under hierarchical models of visual processing, after an initial feedforward sweep of visual information, higher level areas, such as the frontal eye fields, use corollary discharge to provide spatiotemporally precise predictions for lower levels within the visual hierarchy (Schall, Morel, King & Bullier, 1995; Stanton, Bruce & Goldberg, 1995). This ensures that visual stimuli are aligned with their expected post-saccadic retinotopic location despite saccades interrupting the typical flow of visual information. We expect that when the peripheral stimulus entered the first stages of visual processing, information about its current (pre-saccadic) position was combined with information from the CD signal to predict the future (post-saccadic) scene. By activating neuronal receptive fields at the post-saccadic retinotopic location, information can be gathered about the post-saccadic scene to minimise interruptions caused by the saccade.

When the saccade occurred with a longer latency, we instead saw a pattern of decoding very similar to control trials, indicating that the stimulus was being processed at its true retinotopic position (on the opposite side of the screen) rather than the remapped one. The fact that we find no evidence of remapping with longer saccade latencies indicates that the brain must be in a state of motor preparation when the visual information enters the system to elicit such a predictive shift.

The temporal precision afforded by our analysis approach revealed not only the time-window during which stimuli presented before a saccade are remapped, but also when the remapped information about those stimuli is available in the EEG signal. Roughly speaking, remapping occurred for stimuli presented ∼120 ms before saccade onset, but peak decoding was achieved ∼184 ms after stimulus onset (i.e. once the eyes were already in flight). This suggests that predictive remapping as a neural process starts before a saccade, but continues into the trans- and post-saccadic period. It is important to note that the neural response we observed cannot simply be attributed to normal visual processing, since the stimulus was removed from the screen before saccade onset (contrary to some previous studies where the stimulus remained on screen, e.g., Parks & Corballis, 2010; Edwards et al., 2018). This means that there was no direct stimulation of the classical neuronal RFs representing the locations on which the classifiers were trained (i.e., one of the four locations around central fixation). As outlined by Duhamel et al. (1992), the post-saccadic response is reflective of an expectation that the stimulus will fall in the neuronal RF after the eyes arrive at the saccade endpoint. These results are compatible with animal neurophysiological research which found that stimuli presented in close proximity to the saccade were remapped after the saccade (Kusunoki & Goldberg, 2003; Neupane, Guitton & Pack, 2016).

Our results indicate that the timing of remapping is dictated by when sensory information enters the system. This is evidenced by above-chance decoding that is dependent on stimulus presentation timing. When we repeated the same analyses using EEG epochs time-locked to saccade-onset, rather than stimulus onset, there was very little evidence of remapping (Supplementary Figure 1). This is relevant because this supplementary analysis uses exactly the same trials, only realigned in time. However, this temporal alignment to the saccade (rather than the stimulus) causes remapping signals to misalign such that they are smeared in the average. The fact that remapping is locked to the stimulus and not the saccade seems, at first glance, to contradict the idea that remapping is a process driven by a motor action. However, the current analysis exploits the fact that the onset of a visual stimulus triggers waves of activity, both feedforward and feedback (Aggarwal et al., 2022; Dijkstra, Ambrogioni, Vidaurre & van Gerven, 2020; Goddard, Carlson, Dermondy & Woolgar, 2016), that change the activity pattern across the scalp as they propagate through the visual hierarchy (Carlson, Hogendoorn, Kanai, Mesik & Turret, 2011). We trained classifiers using the stimulus evoked activity at a particular timepoint during visual processing. When we time lock the test data to the saccade, visual information will have reached different processing stages in different trials leading to across-trial desynchronisation. This means that the classifiers likely identified remapped stimulus information at different timepoints across trials, which is then lost in the grand average. By time-locking to the stimulus onset, we show that presenting the stimulus during a state of motor planning changes the course of the sensory processing such that it continues along the visual hierarchy as if it were presented at its remapped location.

This interpretation is consistent with findings reported by Edwards et al (2018), who showed that, after saccade landing, it was possible to discriminate a face stimulus from a house stimulus briefly presented in the periphery before a saccade. Contrary to our result, they observed significant above-chance decoding when locking to saccade offset. However, in their study the time between stimulus onset and saccade onset was fixed so that the propagation of feedforward visual information had always reached the same point before the initiation of a saccade. In the present study, the time between stimulus onset and saccade onset was deliberately varied, such that there was a greater variability of stimulus-saccade onset asynchronies (SSOAs) within each saccade bin. If SSOA is variable, and if predictive shifts are driven by the CD signal, then the level at which the CD signal interacts with the visual signal may differ from trial to trial, masking any saccade-locked effects. By locking to the stimulus, we examine how presenting the stimulus in a state of motor preparation transforms its representation in the visual cortex.

Previous studies have argued in favour of attentional shifts as an explanation for saccadic remapping (Cavanagh et al., 2010; Jonikaitis et al., 2013; Melcher & Colby, 2008; Rolfs & Szinte, 2016; Wolf & Schütz, 2015). In the current experiment, both exogenous (abrupt stimulus onset) and endogenous attention (task-relevant stimulus) were cued, meaning attention was engaged on all trials. However, contrary to previous behavioural studies supporting attentional remapping, we did not find remapping when the stimulus was presented significantly before saccade onset. It has been found that presenting an attentional cue considerably before saccade onset (which would correspond to the Middle and Long Bins in the current study), leads to pre-saccadic sensitivity at the remapped location, while presenting the cue close to saccade onset leads to no remapping (Szinte et al., 2018). This suggests possible mechanistic differences between predictive remapping of attention (which has primarily been measured in humans using psychophysical approaches) and spatial remapping of neuronal receptive fields (which has primarily been measured in animal models). Task design may also play a role in this discrepancy. The current study employed a design in which fixation trials, saccade trials and control trials were randomly interleaved, meaning participants could not anticipate when a saccade should be initiated. In contrast, other studies included only saccade trials, meaning that stimulus presentation was always accompanied by a saccade (Szinte et al., 2018). Perhaps, participants used this predictability to prepare a saccade in advance of the saccade cue. This may explain why such studies find remapping despite the attentional cue being presented outside of the normal saccade preparation range (∼200 ms before saccade onset (Fischer & Ramsperger, 1984)). In the current study, attention was always allocated to the stimuli, such that we cannot dissociate between attentional remapping and spatial RF remapping. Further research dissociating attention, for example by including trials in which attention is directed towards a location away from stimulus presentation location, (similar to that implemented while recording from monkey V4; Marino & Mazer, 2018), will therefore be necessary to characterise the role of attention in saccadic remapping.

In conclusion, we provide strong neural evidence in humans of predictive remapping of visual spatial information across saccades. Moreover, we found that the timing of remapping was tightly locked to stimulus presentation, occurred only for stimuli presented in a narrow window immediately preceding a saccade, and continued while the eyes were in motion. Thus, predictive remapping may ensure visual constancy across eye movements by enabling the visual system to represent the pre-saccadic stimulus at its post-saccadic retinotopic position.

## Footnotes

1 Edwards et al. (2016) included one condition in which the stimulus was removed from the screen before the eyes reached the saccade target. This is considered in the discussion section.

