## Supplementary figure 1 for "Decoding Remapped Spatial Information in the Peri-Saccadic Period"

### Supplementary Materials

#### Saccade-locked analysis

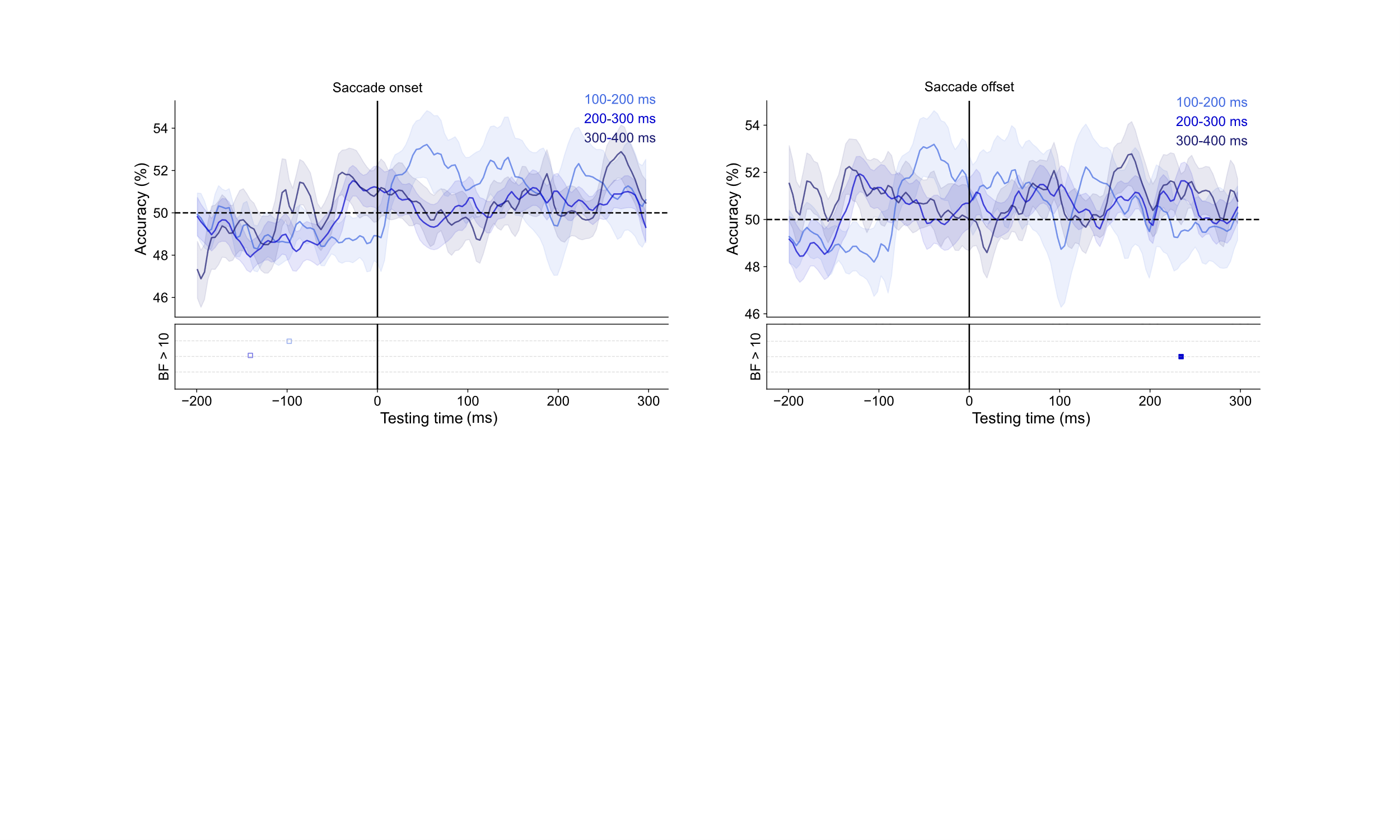
We ran a separate analysis in which the EEG epochs were time locked to the saccade onset and offset, ranging from 200 ms before saccade onset to 300 ms after, in order to investigate whether remapping was dependent on the timing of the saccade, rather than the stimulus. We used the classifiers trained on the peak decoding timepoint for each individual subject (mean = 167 ms) from fixation trials and tested them on EEG data from saccade trials in which stimuli were presented around the saccade target. We examined the time course of saccadic remapping for location information in the time-window from 200 ms before saccade onset/offset to 300 ms after saccade onset/offset. While there seems to be a trend in the Short Bin in which ability to decode the remapped location increases just after saccade onset and just before saccade offset, there was not sufficient evidence to support this result. Locked to saccade onset, below-chance decoding (BF > 10) was found in the Short Bin and the Middle Bin at -98 ms and -141 ms, respectively. Locked to saccade offset, above-chance decoding (BF > 10) was found at 234 ms after saccade offset. However, in all cases evidence was found only at a single timepoint and so is not considered as reliable evidence for a representation of the stimulus in the remapped location. Thus, these results indicate that the evidence for remapping found when locked to stimulus onset is not present when locking to the saccade.

a.

b.

**Supplementary figure 1.** Mean decoding accuracy for saccades occurring at three different latencies relative to stimulus onset (100-200 ms , 200-300 ms, 300-400 ms after stimulus onset), locked to **a.** saccade onset and **b.** saccade offset.  Shaded areas depict standard error of the mean across subjects. Dots below plot indicate thresholded Bayes factors (BF) for which there is substantial evidence for above (filled squares) and below (open squares) chance decoding.
